# Offline knowledge assimilations ensue online obscurity to inspire implicit learning

**DOI:** 10.1101/2021.04.06.438726

**Authors:** Mohamed H. Aly, Kareem Abdou, Reiko Okubo-Suzuki, Masanori Nomoto, Kaoru Inokuchi

## Abstract

Passive priming of prior knowledge to assimilate ongoing experiences underlies advanced cognitive processing. However, the necessary neural dynamics of memory assimilation remains elusive. Uninstructed brain could also show boosted creativity, particularly after idling states, yet it remains unclear whether the idling brain can spontaneously spark relevant knowledge assimilations. We established a paradigm that links/separates context-dependent memories according to geometrical similarities. Mice exploring one of four contexts one day before undergoing contextual fear conditioning in a square context showed a gradual fear transfer to pre-exposed geometrically relevant contexts next day, but not after 15 min. Anterior cingulate cortex neurons representing relevant, rather than distinct, memories were significantly co-reactivated during post-conditioning sleep only, before their selective integration next day during testing. Disrupting sleep co-reactivations prevented assimilation while preserving recent memory consolidation. Thus, assimilating pertinent memories during sleep through co-reactivation of their respective engrams represents the neural underpinnings of sleep-triggered implicit cortical learning.

## Introduction

Flexible information processing that underlies higher-order brain functions, such as applied or implicit learning, decision making, and creative thinking, is thought to rely on the construction of cognitive frameworks^1, 2^. Mnemonic maps structured during spatial learning, for instance, are believed to guide navigation through flexibly imagining unvisited trajectory scenarios while recapturing neural activity sequences of those recently encountered^3–5^. Moreover, implicit ideas, beyond those that are directly trained, are thought to be generated via online assimilation of the ongoing experience with previous knowledge networks that are passively primed with explicit cues or instructions prior to or during learning^6–11^. However, due to the complexity of the implemented learning paradigms and/or technical limitations, causal evidence for the necessity of an interaction between the neural correlate of pre-stored knowledge with that of novel experience for the induction of these knowledge assimilations has not been obtained. Moreover, in such primed learning, assimilations are always initiated online during learning and could even persist throughout later rest^6–11^. Nevertheless, human studies have, on the other hand, documented the ability of idling states such as sleep to spark —without prior priming— the generation of new implicit information^12–16^. However, it remains unclear whether the idling brain machinery compensates for the absence of prior online priming by spontaneously inducing relevant knowledge assimilations to create novel information. In post-encoding sleep, emerging experience-specific reactivations that are thought to be necessary for memory consolidation^17–19^ could be artificially updated to produce a qualitatively novel perception to the acquired experience^20^. We hypothesized that the idling brain triggers the coupling of memory reactivation of relevant existing knowledge with that of recent encounters to enable their flexible assimilation and inspire novel perception. To investigate this hypothesis, here we studied using a simple two-session learning paradigm whether mice can, during sleep alone, bridge temporally distant individual memories that share subtle commonalities to generate, without priming, qualitatively novel information, and how.

Using *in vivo* calcium imaging as well as pharmacological and engram-specific optogenetic manipulations, we unveiled the precise temporal dynamics of multiple memories’ flexible interaction underlying implicit learning. Despite being dispensable for the recent memory consolidation, this interaction —characterized by an increased synchrony among memory engrams— was shown to be crucial for recent memory assimilation to the existing networks of relevant knowledge and for the creation of novel implicit links between potentially related experiences. We further revealed the existence of two different types of memory assimilation, i.e., online and offline, the use of which depended on the degree of overlap between related events. The absence of online assimilations when similarities were inconspicuous during learning was reimbursed in subsequent sleep periods. Thus, offline memory assimilation signifies the necessity of post-learning idling states in compensating for online obscurity, and could therefore explain the previously reported need for sleep to boost creativity^12, 13, 16, 21^. Finally, our results suggest that the dissociation between two parallelly occurring offline processes, namely, recent memory consolidation and knowledge assimilation, was mainly attributed to the division of labor among cortical circuits involving the anterior cingulate cortex (ACC) and prelimbic cortex (PL).

## Results

### Novel experience is assimilated with existing knowledge according to similarities in environmental geometry

Rodents can notice the geometrical similarities and differences between closely and distinctly shaped environments^22^. To study whether mice can assemble context-dependent episodic experiences based on their geometrical commonalities, mice were pre-exposed to one of four different environment conformations – a triangular, hexagonal, octagonal, or circular context (Event 1) (Fig. 1a and Extended Data Fig. 1). One day later, all groups were subjected to contextual fear conditioning (CFC) in a square context (Event 2). On the subsequent day, mice were tested for the transfer of fear memory from the conditioned square to the pre-exposure context (test 1), followed by a second test for the square-associated fear memory 24 h later (test 2). Animals pre-exposed to and tested in the triangular context showed the highest freezing behavior, followed by those subjected to the hexagonal environment (Fig. 1b). Both groups exhibited freezing levels that were not significantly different from those shown in the conditioning context during test 2. On the other hand, octagon or circle pre-exposed mice equally exhibited the least fear memory transfer (Fig. 1b), with significantly lower freezing than that shown in test 2. All groups exhibited equally high freezing in the conditioned square environment (Fig. 1c). This indicates that this effect was driven by the higher degree of geometrical similarity between the triangular, hexagonal, and square environments relative to the circular or octagonal ones.

**Fig. 1.**
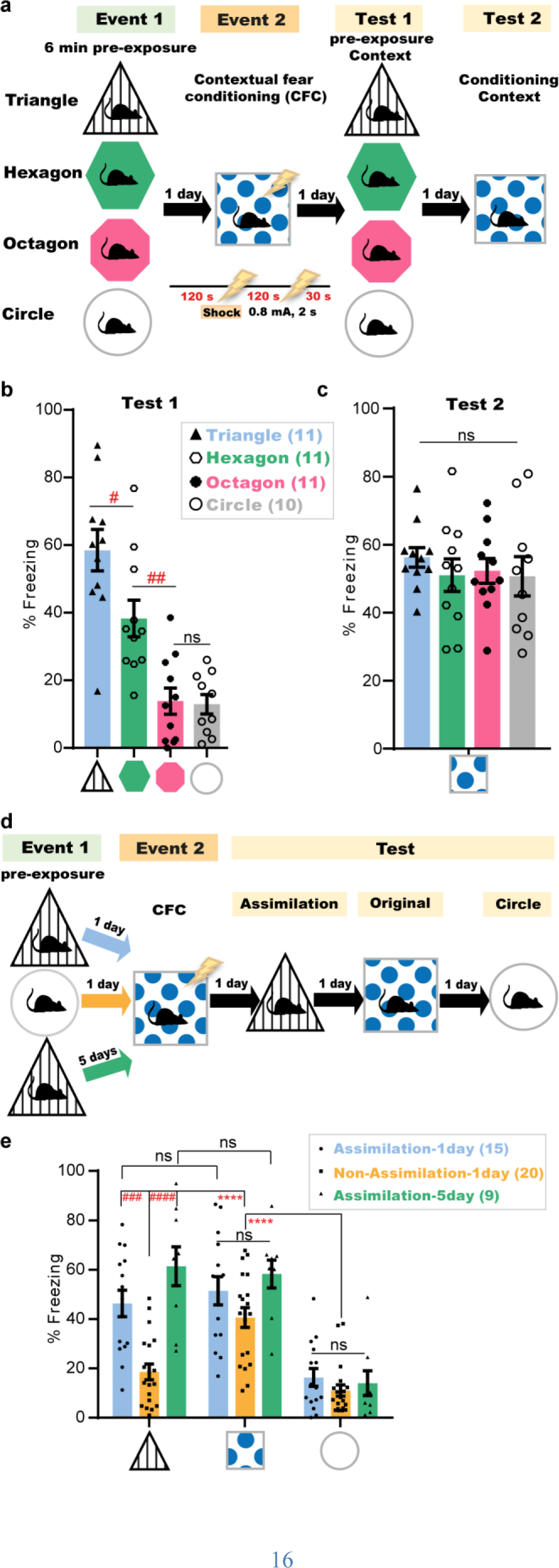
Novel experience is assimilated with existing knowledge according to similarities in environmental geometry. **a,d**, Experimental design; Lightning bolt, electric foot shock. **b,c,** Freezing levels during test (4 min) in the pre-exposure context (**b**), and after 24 h in the conditioned context (**c**). Data are presented as mean ± SEM. *P*-values were determined using a one-way ANOVA followed by Tukey’s multiple comparisons, ^#^*P* < 0.05, ^##^*P* < 0.01. ns, not significant (*P* > 0.05). **e,** Freezing levels during test sessions, 4 min each. Data are presented as mean ± SEM. *P*-values were determined using two-way repeated measures (RM) ANOVA followed by within-group (\*\*\*\**P* < 0.0001) or between-group (^###^*P* < 0.001, ^####^*P* < 0.0001) Tukey’s multiple comparisons. ns, not significant (*P* > 0.05).

It remains unclear whether these contextual similarities are identifiable upon a first-time exposure to a relevant environment. To study this, mice were subjected to two episodic events on two consecutive days (Fig. 1d). In the first event, animals explored either the triangle or circle context. The next day, both groups received two electric foot shocks in the square context and were tested for the fear transfer 24 h later in the triangle context. Mice exposed to the triangle in the first event showed significantly higher freezing than those pre-exposed to the circle context (Fig. 1e). This memory transfer occurred even with a 5-day interval between both events, which indicates its independence on the time of occurrence. Furthermore, all groups exhibited equally high and equally low freezing levels when tested in the square context for the original CFC memory and in the geometrically irrelevant circle context, respectively, on the following 2 days (Fig. 1e). This suggests that the memory transfer, which was specific to related environments, is a process of assimilating two previously acquired relevant experiences rather than a non-specific generalization or an immediate categorization.

### Behavioral assimilation of subtly related experiences occurs only after idling states

To assess whether mice extract these contextual commonalities online during the second event, mice subjected to the triangle environment in Event 1 and to CFC in the square context in Event 2 were tested for assimilation either 15 min after CFC or the next day (Fig. 2a). Animals tested 15 min after CFC had significantly lower freezing levels than those tested one day post-CFC, and lower than their freezing levels in the original CFC memory test (Fig. 2b). However, introducing more obvious commonalities, such as replacing (only during Event 1) the triangle’s spatial cue with that used in the conditioning square (see Extended Data Fig. 1b,c) induced memory transfer even 15 min after CFC. Thus, integration in this primed group was triggered earlier online during the second event, as has been reported by previous studies^7–11^. All groups exhibited equally high and low freezing levels in the original CFC memory and neutral circle tests, respectively (Fig. 2b). These results directly refer to the role of the post-CFC incubation period in the context geometry-based memory assimilation and exclude the possibility of online assimilation during CFC. This incubation-dependent difference in freezing behavior did not exist between the short-term and long-term original CFC memories of the conditioned context (Extended Data Fig. 2). Additionally, even in the absence of an associated valence during Event 2, mice exposed to geometrically relevant environments showed a significant decrease in motility during the assimilation test (Extended Data Fig. 3a,b), which was not simply due to the familiarity with the testing context as a result of repeated exposure^23^. On the other hand, there was no significant difference in the motility between the assimilation and non-assimilation groups during Event 2 (Extended Data Fig. 3c). Taken together, these results further indicate the subtle nature of these commonalities, which mandates offline incubation for the assimilation of these contextual memories to occur.

**Fig. 2.**
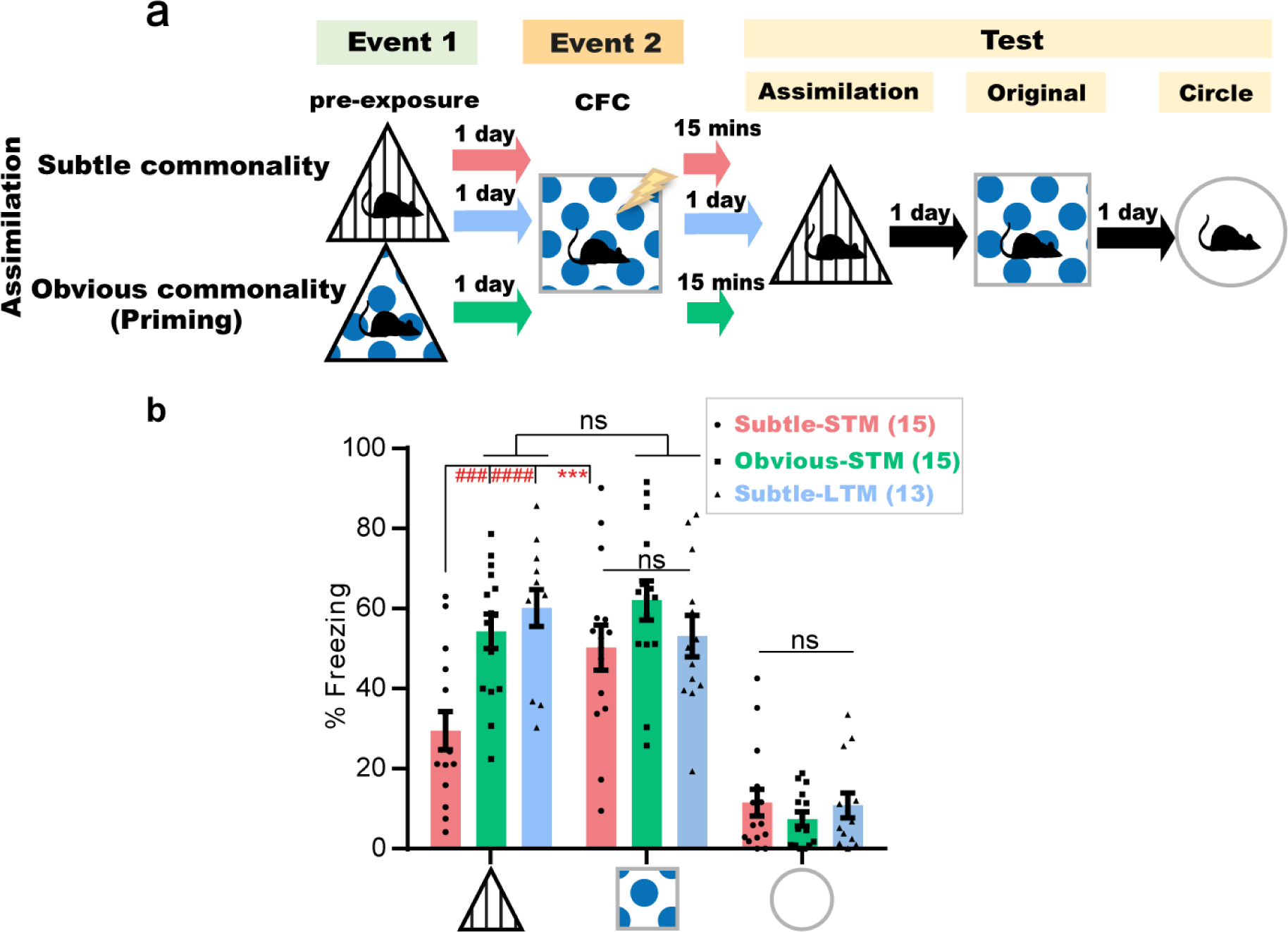
Behavioral assimilation of subtly related experiences occurs only after idling states. **a,** Experimental design. **b,** Freezing levels during test sessions, 4 min each. Data are presented as mean ± SEM. *P*-values were determined using a two-way RM ANOVA followed by within-group (\*\*\**P* < 0.001) and between-group (^####^*P* < 0.0001, ^###^*P* < 0.001) Tukey’s multiple comparisons. ns, not significant (*P* > 0.05). STM, short-term memory; LTM, long-term memory.

**Fig. 3.**
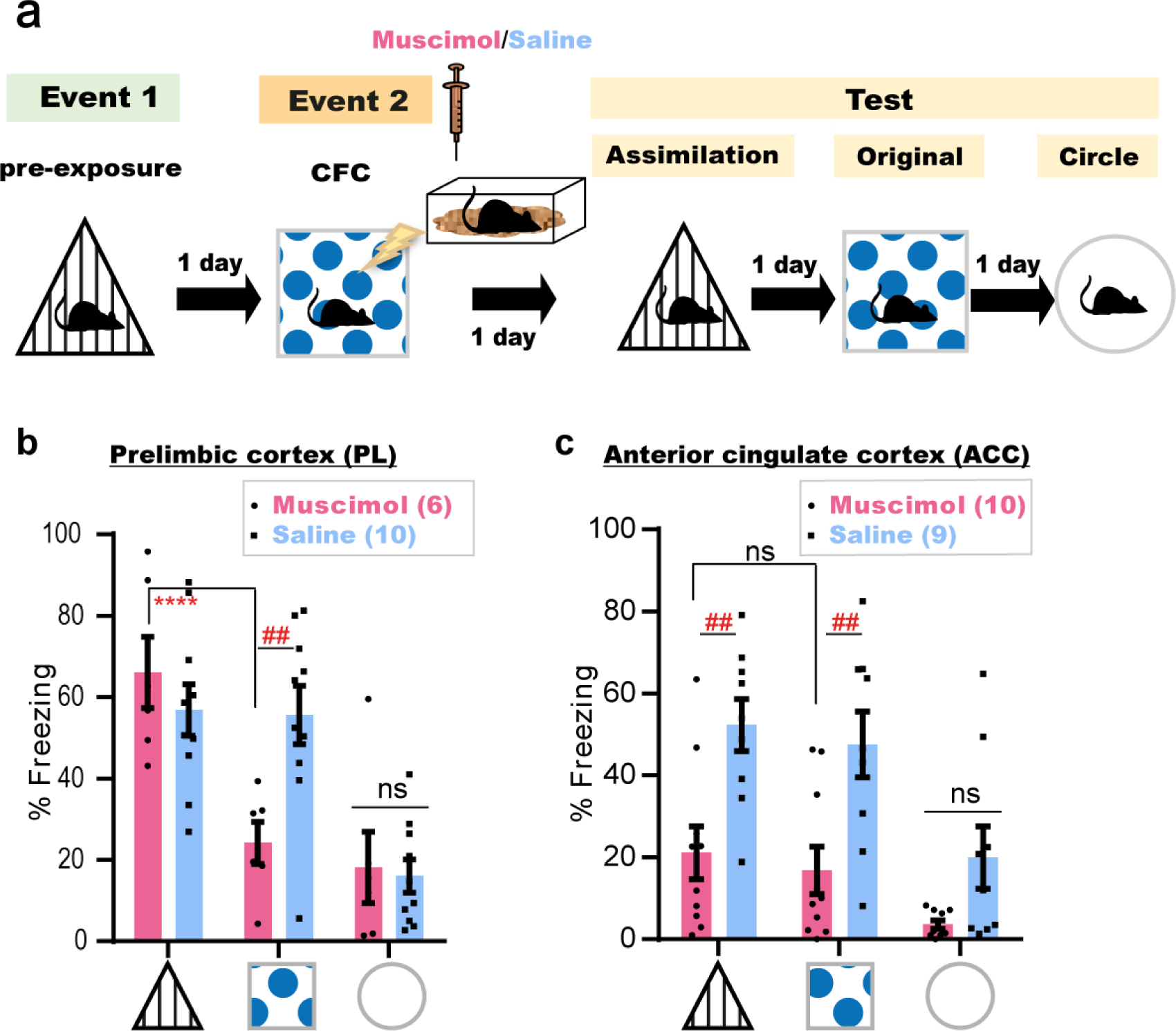
Offline dynamics in the ACC, but not PL, is necessary for memory assimilation. **a,** Experimental design. **b,c,** Freezing levels during test sessions, 4 min each. Data are presented as mean ± SEM. *P*-values were determined using a two-way RM ANOVA followed by within-group (\*\*\*\**P* < 0.0001) Tukey’s and between-group (^##^*P* < 0.01) Bonferroni’s multiple comparisons. ns, not significant (*P* > 0.05).

### Offline dynamics in the ACC, but not PL, is necessary for memory assimilation

The PL and ACC are two established neocortical pillars for the consolidation of associative memory and its reorganization^24–28^. To understand their distinct involvement in each process, mice subjected to the offline assimilation task were bilaterally injected with a γ-aminobutyric acid type A (GABA_A_) receptor agonist, muscimol, into either the PL or ACC immediately after CFC (Fig. 3a and Extended Data Fig.4). Disruption of post-CFC dynamics in the PL did not affect memory assimilation, even though consolidation of Event 2 was compromised (Fig. 3b). However, muscimol injections into the ACC impaired both processes (Fig. 3c). This suggests that post-conditioning ACC dynamics are necessary for the consolidation of a spatial associative memory and its assimilation with stored memories, while offline activity of the PL is mainly concerned with the consolidation of new experiences.

**Fig. 4.**
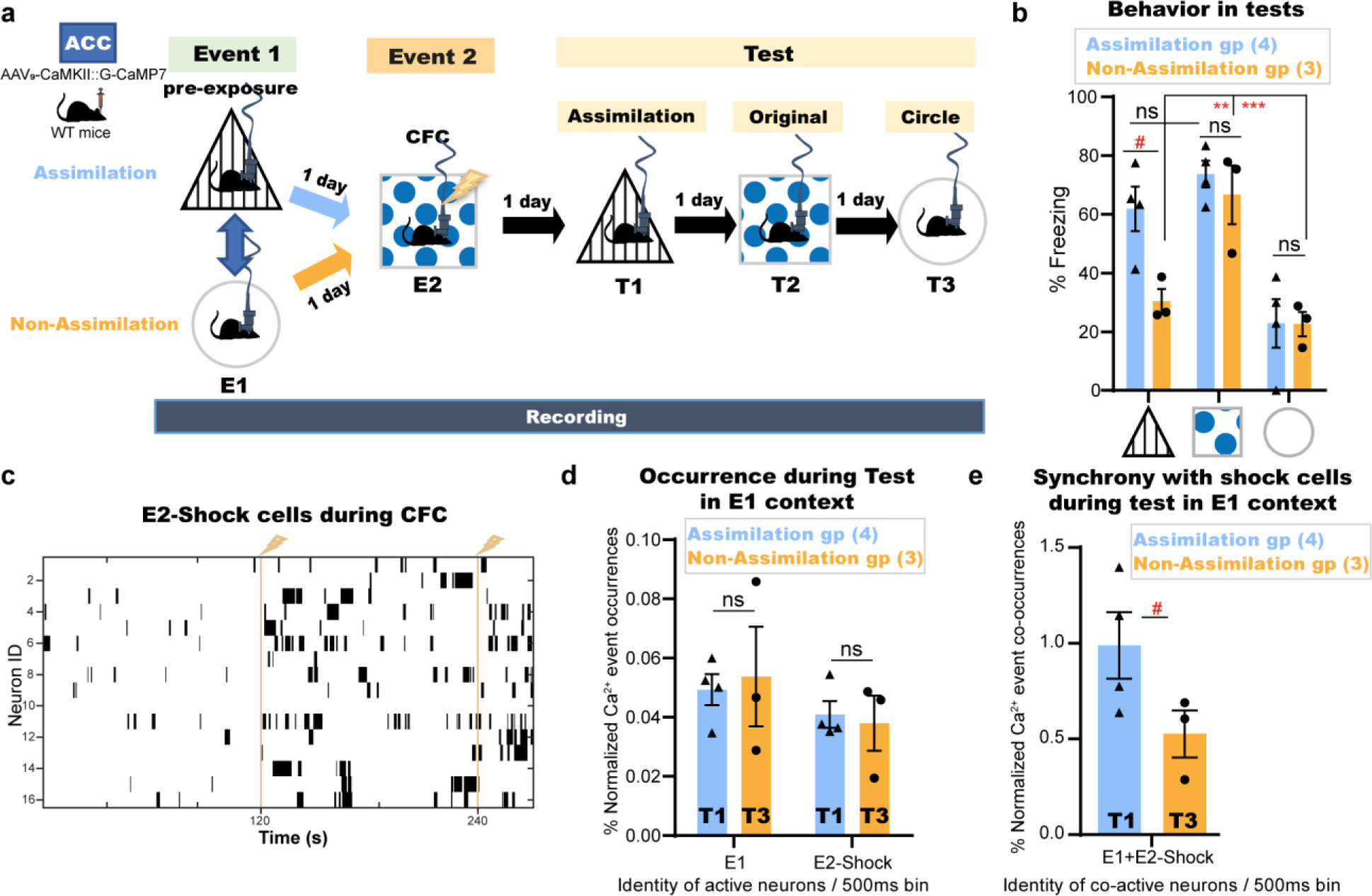
Cells unique to a novel experience are conditionally incorporated into the recall networks of a pre-stored relevant memory in the ACC. **a,** Experimental design. AAV, adeno-associated virus; CaMKII, calcium/calmodulin-dependent protein kinase II; WT, wild type; E1, event 1; E2, event 2; T1, test 1; T2, test 2; T3, test 3. **b,** Manually calculated freezing levels during test sessions (4 min each). **c,** Example Ca^2+^ transients for E2-Shock cells during CFC; Vertical yellow lines, shock onset. **d,** Single Ca^2+^ event occurrences of E1-cells or E2-Shock cells during the respective recall sessions (Assimilation, T1; Non-Assimilation, T3) of E1 pre-exposure context. **e,** Synchronized Ca^2+^ event co-occurrences of E1-cells and E2-Shock cells during the respective recall sessions (Assimilation, T1; Non-Assimilation, T3) of E1 pre-exposure context. Data are presented as mean ± SEM in **(b, d, and e)**. *P*-values were determined using a two-way RM ANOVA followed by within-group Tukey’s tests or between-group Bonferroni’s multiple comparisons **(b)** or using unpaired *t* test **(d and e)**. For within-group comparisons (\*\*\**P* < 0.001, \*\**P* < 0.01); For between-group comparisons (^#^*P* < 0.05). ns, not significant (*P* > 0.05).

### Cells unique to a novel experience are conditionally incorporated into the recall networks of a pre-stored relevant memory in the ACC

To further investigate how assimilation is processed in the ACC, mice were injected with Adeno-Associated Virus 9 (AAV_9_) encoding calcium/calmodulin-dependent protein kinase II (CaMKII)::G-CaMP7 into the ACC (Extended Data Fig. 5a) to record the underlying dynamics of this task using a head-mounted miniature fluorescent microscope^29^. The same ACC neurons were accurately tracked across sessions^30^ along the offline memory assimilation task (Fig. 4a,b). An automated sorting system (HOTARU)^31^ was used to extract each neuron’s calcium (Ca^2+^) activity throughout the task (see methods for details). Subsequently, Ca^2+^ event frequency of every neuron during each event was automatically calculated, and accordingly, each neuron was classified as Event 1 (E1)-, Event 2 (E2)-Square-, or E2-Shock-responsive cells based on their differential Ca^2+^ event frequencies (Fig. 4c and Extended Data Fig. 5b; see methods for details). The occurrences of Ca^2+^ events for neurons representing each class were automatically enumerated per 500 ms in each session (Fig. 4d and Extended Data Fig. 5e). The synchronization of Ca^2+^ event occurrences of E2-Shock neurons with those of E1-responsive cells was significantly higher in the assimilation group relative to the non-assimilation group during the respective recall session of the pre-exposure context (Fig. 4e). However, there were no significant differences in single Ca^2+^ event occurrences of each neuron category between the assimilation and non-assimilation groups during these recall sessions (Fig. 4d).

**Fig. 5.**
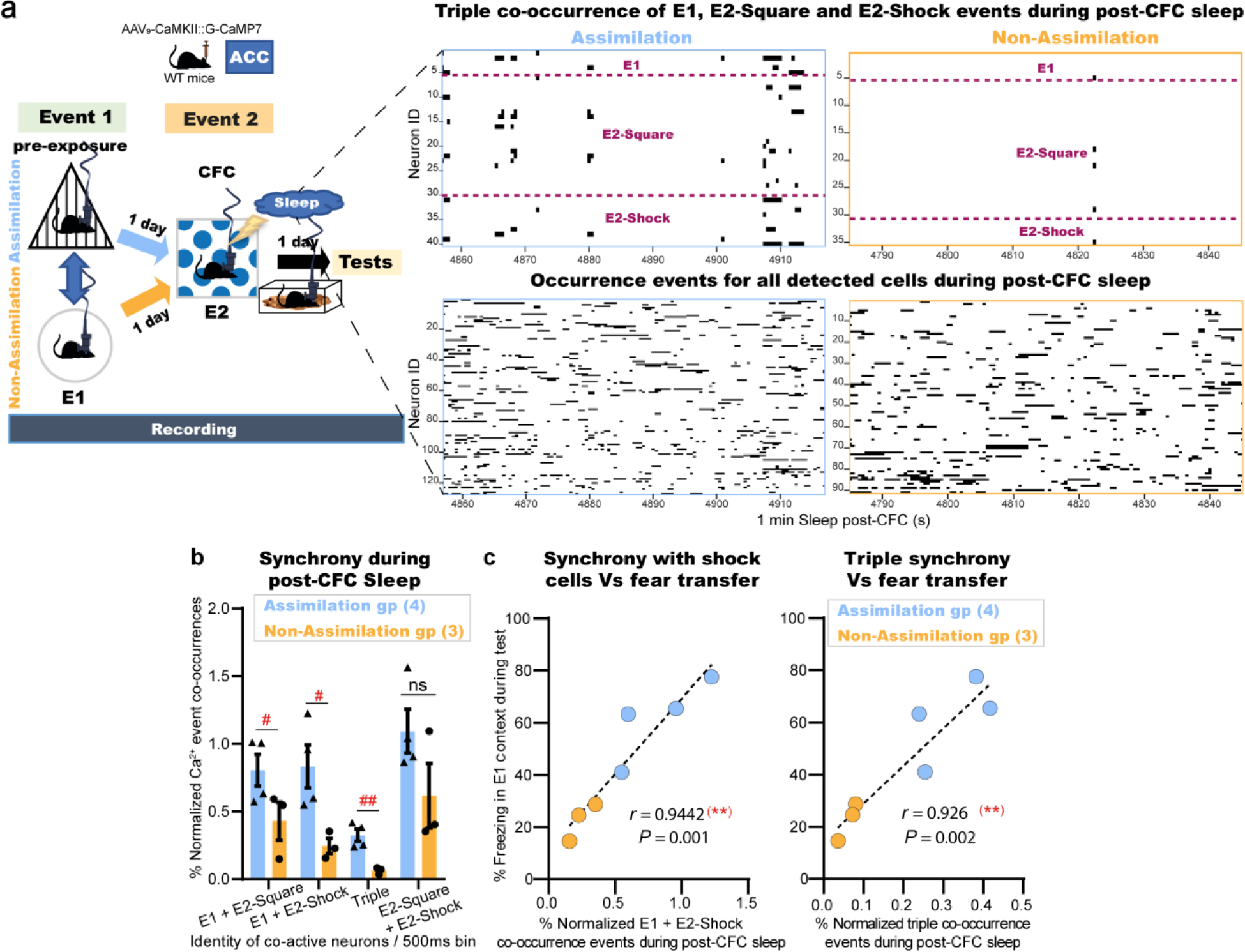
Co-reactivations of related memory representations in the ACC emerge during post-learning sleep. **a,** Left: Experimental design. Right: Representative plots for 1 min of post-CFC sleep in the assimilation (blue) and non-assimilation (yellow) groups depicting either the synchronized Ca^2+^ transients for the triple co-occurrence events of E1, E2-Shock, and E2-Square neurons (top), or the occurrence events of all detected neurons (bottom). **b,** Percent normalized Ca^2+^ event co-occurrences of either E1-responsive cells with E2-responsive cells or E2-Square with E2-Shock responsive cells within 2 min post-CFC sleep. Data are presented as mean ± SEM. *P*-values were determined using unpaired *t* test; (^##^*P* < 0.01, ^#^*P* < 0.05). ns, not significant (*P* > 0.05). **c,** Individual percent freezing during test in the E1 context, as in Fig. 4b, and the synchronized Ca^2+^ events of E1-cells with that of E2-Shock cells alone (left) or with that of both E2-Square and E2-Shock cells (right) during sleep, as in **(b)**; Dashed black line, linear fit; Inset, Pearson’s (r) value and *P*-value (two-tailed).

### Co-reactivations of related memory representations in the ACC emerge during post-learning sleep

During sleep periods after the second event, there were significantly more synchronized Ca^2+^ event occurrences of E1-responsive neurons with those of E2-responsive cells in the assimilation group than in the non-assimilation group (Fig. 5a,b and Extended Data Fig. 5d). On the other hand, there was no significant difference between the two groups in the Ca^2+^ event co-occurrences of E2-Square with E2-Shock cells representing the association that occurred earlier during this event (Fig. 5b). Moreover, there was a positive correlation between the synchronized event co-occurrences of E1-cells and E2-cells during post-CFC sleep in each group and the freezing behavior during the recall session of the pre-exposure context (Figs. 5c and 4b). By contrast, event co-occurrences of E1 and E2 neurons during CFC training were not significantly different between the two groups (Extended Data Fig. 5c). Finally, there were no significant differences in single Ca^2+^ event occurrences of each neuron category between the assimilation and non-assimilation groups during either session (Extended Data Fig. 5e). These results strongly suggest that the emergence of neuronal co-activity during post-CFC sleep was dependent on the presence of relevance between both events. The correlation of this offline co-activity with the later fear transfer in addition to its earlier absence throughout E2 are in line with our behavioral results (Fig. 2a,b and Extended Data Fig. 3), both referring to the offline initiation of this process.

### Interaction of pre-stored memories with new related experiences during sleep is necessary to induce offline assimilation

To examine the importance of co-activities during sleep in memory assimilation, a combination of two AAVs encoding activity-regulated E-SARE::ER^T2^CreER^T2^-PEST and CaMKII::DIO(loxP)-eArch3.0-eYFP were injected into the ACC of the mouse brain to label the engram of Event 1 with a light-sensitive neural silencer in a tamoxifen-inducible manner (Fig. 6a,b; see methods for details). Subsequently, mice were intraperitoneally injected either with 4-Hydroxytamoxifen (4-OHT) or vehicle 2 h after triangle pre-exposure and were subjected to CFC five days later in the square context (Fig. 6b). A continuous laser (589 nm) was bilaterally delivered to the ACC during any subsequent sleep within 3-4 h of the post-CFC period (Fig. 6c). In the assimilation test the next day, 4-OHT-injected animals showed significantly lower freezing behaviors than that exhibited in the conditioned context test performed one day later (Fig. 6d, Left). However, in the absence of either 4-OHT or offline light stimulation, mice showed successful assimilation. On the other hand, there were no significant differences in the freezing levels between the groups in the original CFC memory test, nor in their cumulative sleep stage-specific laser illumination duration (Fig. 6d, Right). Thus, the spontaneous synchronized reactivation of relevant stored engrams is necessary for new memory networking and the resultant implicit learning to occur, but dispensable for its consolidation.

**Fig. 6.**
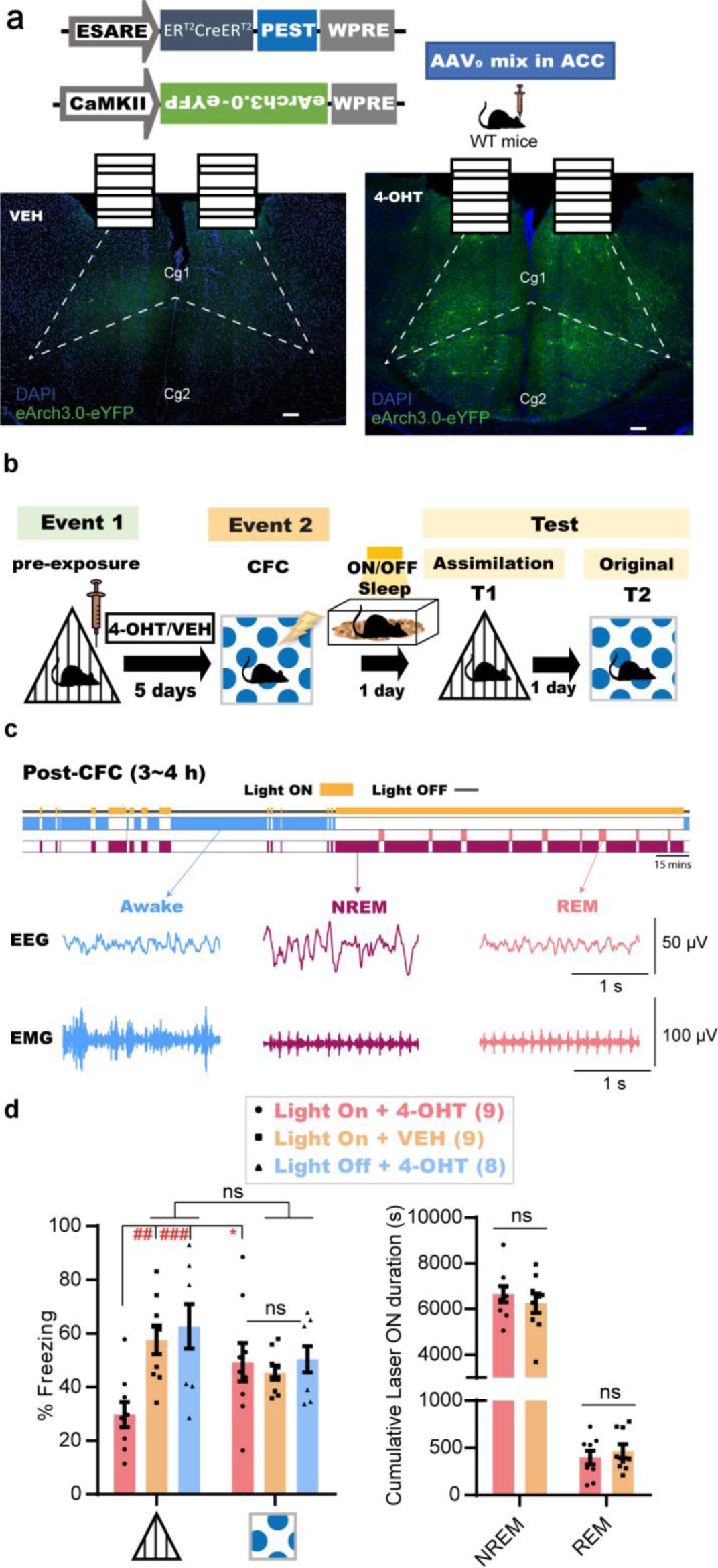
Interaction of pre-stored memories with new related experiences during sleep is necessary to induce offline assimilation. **a,** Top: Design of AAVs injected into the ACC for engram labeling. Bottom: Coronal section of the ACC showing eYFP protein expression with VEH (left) or 4-OHT (right). Dashed lines show the boundary of the ACC. Scale bar represents 100 μm. ER^T2^CreER^T2^, tamoxifen-inducible recombinase; eYFP, enhanced yellow fluorescent protein; DAPI, 4՛,6-diamidino-2-phenylindole; Cg1, cingulate cortex, area 1; Cg2, cingulate cortex, area 2; 4-OHT, 4-hydroxytamoxifen; VEH, vehicle. **b,** Experimental design. ON; laser ON group; OFF, laser OFF group. **c,** Top: Diagram for sleep-specific optogenetics. Bottom: Example EEG and EMG recordings for each detected stage. EEG, electroencephalogram; EMG, electromyogram; NREM, Non-rapid eye movement; REM, rapid eye movement. **d,** Left: Freezing levels during test sessions. Graph shows mean ± SEM. *P*-values were determined using a two-way RM ANOVA followed by within-group Bonferroni’s (\**P* < 0.05) or between-groups Tukey’s (^##^*P* < 0.01, ^###^*P* < 0.001) multiple comparisons. ns, not significant (*P* > 0.05). Right: Cumulative duration of laser illumination in each sleep stage across groups. Graph shows mean ± SEM. *P*-values were determined using a two-way RM ANOVA followed by between-groups Sidak’s (ns, not significant (*P* > 0.05) multiple comparisons.

## Discussion

Our findings demonstrate the necessity of compensatory idling brain activities to uncover commonalities between temporally distant yet related experiences, in the absence of explicit online cueing for their relevance, thus triggering their integration. The calcium imaging results strongly suggest that there is a conditional incorporation of cells unique to a novel experience into the recall networks of a pre-stored relevant memory during the later expression of their integration. The inability of mice to behaviorally extract these similarities online during learning, due to the lack of obvious cues, directly demonstrates the involvement of post-encoding periods in such reorganization of neural networks. Spontaneous co-reactivation of ACC neuronal representations of these subtle commonality-bearing experiences, which was strongly correlated with their future integration, emerged during post-encoding sleep rather than throughout the learning. Disrupting relevant memories’ integration by inhibiting these synchronized offline reactivations of the pre-stored ACC engram cells establishes the necessity of these co-reactivations as a physiological mechanism for restructuring the existing memory networks to assimilate novel related experience.

However, the elimination of online obscurity during learning by exposure to more obvious commonality-bearing experiences spared the need for these compensatory post-encoding offline processing to initiate the process of integration, which seems to have begun earlier online during the second experience. These results are also in line with previous studies, in which the priming of prior knowledge with conspicuous cues or explicit instructions online during a novel experience learning led to their immediate integration^7–11^. These studies mainly applied a sensory pre-conditioning task, which implements a shared obvious element (cue) between two learning experiences. Integration of both experiences in this task has been shown to occur online^9–11^ due to the co-reactivation of hippocampal representations of the first event during exposure to the second^9^. Moreover, using MEG scanning for human subjects performing an instruction-based applied learning task, another study observed the online emergence of synchronized representations from two overlapping episodes during the successful engagement in the second one^8^. Our results could further demonstrate the necessity of such synchrony for triggering the integration of relevant experiences. Additionally, we further revealed that, in the absence of prior online priming, compensatory post-learning sleep co-reactivations of cortical (ACC) representations of subtly related experiences emerge to inspire their implicit connections by creating novel associations. The co-activations reported in our study could be different from those observed in a recent study that implemented sensory pre-conditioning task in humans and mice^7^. In that recent study, the offline co-activations of hippocampal representations of conspicuously overlapping experiences did not appear until days after their successful behavioral association, and therefore could have been just reflecting the reconsolidation of the already learned association rather than triggering it. Moreover, considering a potential impact of the randomized co-incident retraining employed in that study on the successive testing days, these co-activations could have been reinstating the memory-chaining representations of an online-triggered second-order conditioning^32^ or those of a repeated co-retrieval^11^. Nevertheless, the role of the hippocampus in either orchestrating or reconsolidating the compensatory offline cortical learning reported in our study remains unknown and further research should investigate the possible involvement of a hippocampal-cortical dialogue during the different stages of this offline memory assimilation task.

Interestingly, non-specific pharmacological disruption of the post-learning dynamics in the ACC impaired both recent experience consolidation and assimilation processes, while similar perturbations in the PL only affected consolidation. Additionally, the selective impairment of recent memory assimilation while sparing its consolidation using ACC engram-specific optogenetic manipulations further ascertained the dual role of the ACC during post-learning sleep. These results also reveal the division of offline memory processing labor among cortical circuits, thereby permitting the separation of a new episode processing and its assimilation. However, despite the dissociation between both processes in a higher-order cognitive function such as in implicit learning, this relationship could differ in the case of applied or schema-based learning in which the consolidation of a new experience might be conditional on its initial assimilation to previously established networks^2, 8^. Such a discrepancy could reflect two opposite directions of memories’ interaction i.e., a new event assimilating to old networks and vice versa.

Online co-reactivation, either physiologically or artificially, of engram cells representing distinct memories generates associative memories, in which a co-reactivation-induced increase in the sharing of engram cell populations mediates the association^33, 34^. Some engram cell ensembles (sub-ensembles) are selectively reactivated in post-learning sleep. These sub-ensembles are preferentially stabilized and reused in retrieval, thus serving as a memory trace^30^. A similar mechanism of selective co-reactivation and sharing of engram cells operates during post-learning sleep to extract a specific commonality for the assimilation of a new event with relevant memory networks. Such offline-exclusive cortical learning mechanism could be the neural underpinnings of sleep-triggered creativity and insight that lead to the inference of novel ideas necessary for survival and adaptive decision making^16^.

## Methods

### Subjects

Naïve wild-type male C57BL/6J mice were purchased from Sankyo Labo Service Co. Inc. (Tokyo, Japan) and maintained on a 12-h light/dark cycle at a controlled temperature (24°C ± 3°C) and humidity (55% ± 5%) with free access to food and water. Mice used in behavioral experiments were 12–18 weeks old. All experimental procedures with animals were congruent with the guidelines of the National Institutes of Health (NIH). The Animal Care and Use Committee of the University of Toyama approved all animal proceedings.

### Viral constructs

For the *in vivo* Ca^2+^ imaging experiment, the recombinant Adeno-Associated Virus (AAV) vector used was AAV_9_-CaMKII::G-CaMP7 (9.4 × 10^12^ vg/mL)^35^ after 10-fold dilution with phosphate buffered saline (PBS) (T900, Takara BIO INC, Japan) before injection. The pAAV-CaMKII-G-CaMP7 plasmid was previously constructed^35^. For tamoxifen-inducible engram labeling^36–38^ (Fig. 6a,b) in the optogenetics experiment^38^, the combination of AAV vectors used was AAV_9_-E-SARE::ER^T2^CreER^T2^-PEST (9.01 × 10^13^ vg/mL) and AAV_9_-CaMKII::DIO(loxP)-eArch3.0-eYFP (6.55 × 10^14^ vg/mL) at a 1:1 ratio. For the pAAV-E-SARE::ER^T2^CreER^T2^-PEST preparation, pAAV-E-SARE::DIO(loxP)-hKikGR was first constructed by replacing the PCR-amplified hKikGR of pLenti-CaMKII::Chrimson-TS-P2A-hKikGR (kindly donated by Dr N. Ohkawa) with primers (sense, TTTTCAATTGGCCACCATGGTGAGCGTGATC; antisense, TTTTTGTACATTACTTGGCCAGCCTGGGCAG) into pAAV-E-SARE::DIO(loxP)-eArchT3.0-eYFP (described previously^34^) at the *BsrG*I-*Mfe*I restriction sites. Then, the *Sma*I-*Ecor*I-digested ER^T2^CreER^T2^-PEST synthesized sequence (Eurofin Genomics Co. Ltd., Tokyo, Japan) was subcloned into the *Hinc*II-*Ecor*I restriction site of pAAV-E-SARE::DIO(loxP)-hKikGR, generating pAAV-E-SARE::ER^T2^CreER^T2^-PEST. Construction of pAAV-CaMKIIα::DIO(loxP)-eArch3.0-eYFP was done by replacing the EF1α fragment at the *Mul*I-*Sal*I restriction site of pAAV-EF1α::DIO(loxP)-eArch3.0-eYFP (kindly donated by Dr K. Deisseroth) with a CaMKIIα fragment of pAAV-CaMKIIα-hM4Dq(Gi)-mCherry (Addgene plasmid 50477). Recombinant AAV_9_ production was performed using the Minimal Purification (MP) method, and the viral genomic titer was subsequently calculated as described previously^39^. Briefly, pAAV recombinant vectors were produced using HEK293 T cells (AAV293; 240073, Agilent Tech, CA, USA) cultured in 15-cm dishes (Corning, NY, USA). Cultured cells were maintained in Dulbecco’s Modified Eagle Medium (D-MEM) (11995-065, GIBCO life technologies, USA) supplemented with 10% fetal bovine serum (FBS) (10270106, GIBCO life technologies, USA), 1% 2 mM L-Glutamine (25030-149, GIBCO life technologies, USA), 1% 10 mM non-essential amino acid (MEM NEAA 100x, 11140-050, GIBCO life technologies, USA), and 1% (100x) penicillin-streptomycin solution (15140-148, GIBCO life technologies, USA). Confluent (70%) HEK293 T cells were transfected using medium containing the constructed expression vector, pRep/Cap, and pHelper (240071, Agilent Technologies, Santa Clara, CA, USA) mixed with the transfection reagent polyethylenimine (PEI) hydrochloride (PEI Max, 24765-1, Polysciences, Inc., Warrington, PA, USA) at a 1:2 ratio (W/V). After 24 h, the transfection medium was discarded, and cells were incubated for another 5 days in an FBS-free maintenance medium. On day 6, the AAV-containing medium was collected and purified from cell debris using a 0.45 μm Millex-HV syringe filter (SLHV033RS, Merck Millipore, Germany). The filtered medium was concentrated and diluted with D-PBS (14190-144, GIBCO Life Technologies, USA) twice using the Vivaspin 20 column (VS2041, Sartorius, Germany) after blocking the column membrane with 1% bovine serum albumin (01862-87, Nacalai Tesque, Inc., Japan) in PBS. To further calculate the titer, degradation of any residual cDNA in the viral solution from production was first assured by Benzonase nuclease treatment (70746, Merck Millipore, Germany). Subsequently, viral genomic DNA was digestion with proteinase K (162-22751, FUJIFILM Wako Pure Chemical, Osaka, Japan) and extraction with phenol/chloroform/isoamyl alcohol 25:24:1 v/v, followed by precipitation with isopropanol and final dissolution in TE buffer (10 mM Tris [pH 8.0], 1 mM EDTA). Titer quantification for each viral solution, referenced to that of the corresponding expression plasmid, was done by real-time quantitative PCR (qPCR) using THUDERBIRD SYBR qPCR Master Mix (QRS-201, Toyobo Co. Ltd., Japan) with the primers 5′ - GGAACCCCTAGTGATGGAGTT-3′ and 5′-CGGCCTCAGTGAGCGA-3′ targeting the ITR sequence. The cycling parameters were adjusted as follows: initial denaturation at 95°C for 60 s, followed by 40 cycles of 95°C for 15 s and 60°C for 30 s.

### Drugs

Muscimol (M1523, Sigma-Aldrich) was dissolved in saline and divided into single-use aliquots before storage at −80°C. 4-Hydroxytamoxifen (4-OHT) (H6278, Sigma-Aldrich) was handled as previously described^37, 40^. Briefly, 4-OHT (10 mg) was dissolved in 250 μl dimethyl sulfoxide (DMSO) (049-07213, FUJIFILM Wako Pure Chemical, Osaka, Japan) and aliquoted into single-use 50 μl solutions before storage in −80°C. Each freshly thawed aliquot was mixed with 1 ml saline containing 2% Tween 80 (P1754, Sigma-Aldrich) and then diluted with 1 ml saline to reach the final I.P. injectable concentration of 1 mg/ml 4-OHT in 2 ml vehicle containing 2.5% DMSO and 1% TWEEN80 solution in saline.

### Surgery

Mice (12–14 weeks old) were given an intraperitoneal anesthesia injection^35^ containing 0.75 mg/kg medetomidine (Domitor; Nippon Zenyaku Kogyo Co. Ltd., Japan), 4.0 mg/kg midazolam (Fuji Pharma Co. Ltd., Japan), and 5.0 mg/kg butorphanol (Vetorphale, Meiji Seika Pharma Co. Ltd., Japan) before being placed, after sedation, on a stereotactic apparatus (Narishige, Tokyo, Japan). After surgery, an intramuscular injection of 1.5 mg/kg atipamezole (Antisedan; Nippon Zenyaku Kogyo Co., Japan), an antagonist of medetomidine, was administered to boost recovery from sedation. Mice were allowed to recover from surgery for 3 weeks in their home cages before behavioral experiments were initiated.

For pharmacological experiments, a double guide cannula (C2002GS-5-0.7/SPC, diameter 0.29 mm, Plastics One Inc., USA) composed of two 0.7 mm-spaced stainless steel pipes protruding 2 mm from the plastic cannula body and fitted with a double injection cannula protruding a further 0.5 mm (C2002IS-5-0.7/SPC, diameter 0.125 mm, Plastics One Inc., USA) was bilaterally inserted either into the anterior cingulate cortex (ACC) (from bregma: +1.0 mm anteroposterior [AP], ±0.35 mm mediolateral [ML], from the skull surface: +1.5 mm dorsoventral [DV]) or the prelimbic cortex (PL) (from bregma: +2.0 mm AP, ±0.35 mm ML, from the skull surface: +1.8 mm DV). Guide cannulas were fixed using dental cement (Provinice, Shofu Inc., Japan) into micro-screws that were anchored into the skull near bregma and lambda. After complete fixation, the injection cannula was removed and replaced by a dummy cannula (C2002DCS-5-0.7/SPC, protrusion 0 mm, Plastics One Inc., USA) to protect from particulate matter.

All virus injections were done using a 10 µl Hamilton syringe (80030, Hamilton, USA) that was fitted with a mineral oil-filled glass needle and wired to an automated motorized microinjector IMS-20 (Narishige, Japan). For the optogenetics experiment, 1000 nL of AAVs mix was injected at 200 nL min^−1^ bilaterally into the ACC (from bregma: +1.0 mm AP, ±0.35 mm ML, from the skull surface: +1.5 mm DV). The glass injection tip was maintained after injection at target coordinates for an additional 5 min before being removed. After 3 weeks of recovery from AAV injection surgery, re-anesthetized mice were placed once again on a stereotactic apparatus to insert, using a manipulator (Narishige, Japan), a stainless guide cannula (C2002GS-5-0.7/SPC, diameter 0.29 mm, Plastics One Inc., USA) 1.0 mm ventral to the skull surface at the ACC coordinates. A guide cannula was then anchored onto the skull with dental cement and a dummy cannula (C2002DCS-5-0.7/SPC, protrusion 0 mm, Plastics One Inc., USA) was subsequently inserted into the guide cannula for protection. In parallel, a custom-built electroencephalogram/electromyography (EEG/EMG) 5-pin system was installed into the skull, as previously described ^41^. Briefly, electrodes were screwed into the parietal cortex for EEG recordings, into the right cerebellar cortex for grounding, and into the left cerebellar cortex as a reference. Additionally, two wires were implanted in the neck muscle for EMG recording. Finally, dental cement was used to fix all system screws in place.

For the Ca^2+^ imaging experiment, 500 nL of AAV_9_-CaMKII::G-CaMP7 was injected at 100 nL min^−1^ unilaterally into the left ACC (+1.0 mm AP, +0.35 mm ML, +1.5 mm DV). After 2 weeks of recovery from AAV injection surgery, re-anesthetized mice were placed once again on a stereotactic apparatus to implant a gradient index (GRIN) lens^29, 35, 42^ (0.5 mm diameter, 4 mm length; Inscopix Inc., USA) into the center of injection (from the skull surface: +1.2 mm DV) using custom-made forceps attached to a manipulator (Narishige, Japan). A low-temperature cautery was used to emulsify bone wax into the gaps between the GRIN lens and the skull, and then the lens was anchored in place using dental cement. Additionally, a custom-built EEG/EMG 5-pin system was installed and cemented into the skull, as mentioned earlier. Three weeks after GRIN implantation, mice were re-anesthetized and placed back onto the stereotactic apparatus to set a baseplate (Inscopix Inc.), as described previously^29, 30, 35, 42^. In brief, a Gripper (Inscopix Inc.) holding a baseplate attached to a miniature microscope^29^ (nVista HD v3; Inscopix, Inc.) was lowered over the previously set GRIN lens until visualization of clear vasculature was possible, indicating the optimum focal plane. Dental cement was then applied to fix the baseplate in position to preserve the optimal focal plane. Mice recovered from surgery in their home cages for 1 week before behavioral imaging experiments began.

### Offline memory assimilation task

Mice in their home cages were transferred using carts during the light cycle from racks in the maintenance room to a similar rack in a soundproof waiting room (A) (Yamaha Co., Shizuoka, Japan). Mice were then left undisturbed for at least 15 min before behavioral experiments began. All behavioral sessions were conducted in another soundproof behavioral room (B) (Yamaha Co., Shizuoka, Japan). For the *in vivo* Ca^2+^ imaging experiment (Figs. 4, 5, and Extended Data Fig. 5), a head-mounted nVista HD microscope was fitted once the animal arrived in waiting room A to adjust the focal plane, and it was kept attached to the subject during transport to behavioral room B. At the end of each session, animals were transported back to the waiting room A and were kept undisturbed for at least 15 min before being moved back in their home cages using carts to their position on the racks in the maintenance room. The assimilation task was composed of two learning sessions (Events 1 and 2) followed by three testing sessions, as follows:

Event 1 (Pre-exposure): At the start of this session, one mouse in its home cage was moved from waiting room A into well-lit behavioral room B and then placed in a pre-exposure context (see Extended Data Fig. 1a,b,d-f). The mouse explored this context freely for 6 min before being removed, returned to its home cage, and moved back to waiting room A. Contexts were cleaned using water and 70% ethanol after each subject. For engram labeling experiments (Fig. 6a,b), mice were injected with 4-OHT (I.P., 10 mg/kg) after being subjected to isoflurane (099-06571, FUJIFILM Wako Pure Chemical, Osaka, Japan) anesthesia 2–3 h after Event 1.

Event 2: One or five days later, each individual mouse was moved blindly from waiting room A in a well-covered and adequately aeriated transport cage to the darkened behavioral room B. The mouse was then subjected to contextual fear conditioning (CFC) after being placed in a square context (see Extended Data Fig. 1c) with dim lighting, benzaldehyde odor (originating from a dish containing 3 cotton buds soaked in 0.25% benzaldehyde/water), and 26 stainless steel floor rods (diameter 2 mm, 5 mm apart) connected to a shock generator (SGS-003, MUROMACHI KIKAI Co. Ltd. Japan) via a cable. Briefly, the subject freely explored the context for 2 min before receiving two electric foot shocks (2 s, 0.8 mA) with a 2-min interval, and was removed 30 s after the second shock. For the motility experiment (see Extended Data Fig. 3), mice in Event 2 explored either the square context (dim light conditions) or the circle context (well-lit conditions) (see Extended Data Fig. 1c,d) for 4.5 mins without being subjected to the electric foot shocks. At the end of the session, animals were returned blindly in the covered transport cage to their home cage in waiting room A. Contexts were cleaned using water and 70% ethanol after each subject.

For pharmacological manipulation experiments (Fig. 3 and Extended Data Fig. 4), 1 μl muscimol (0.625 μg/μl) or an equivalent volume of vehicle (saline) was injected per hemisphere into the PL or ACC at a flow rate of 0.2 µL/min under isoflurane anesthesia once the animal had been moved back to waiting room A, directly after the CFC. Intracranial injections were performed using the previously mentioned double injection cannula connected with two (10 μl) Hamilton syringes (84853, Hamilton, USA) placed in a micro-syringe pump (CMA 400, Harvard Apparatus). The injection cannula was left for 5 min after injection to allow for complete drug diffusion before being removed and replaced with the dummy cannula mentioned earlier. Injections were repeated one more time after 2.5 h to extend the drug effect.

For the *in vivo* Ca^2+^ imaging experiments (Figs. 4, 5, and Extended Data Fig. 5), each mouse after CFC training, still attached to the nVista HD microscope in waiting room A, was moved in its home cage back to rather a well-lit behavioral room B, placed in a sleep box, and simultaneously connected to the EEG/EMG recording unit. To avoid photobleaching, the nVista light-emitting diode (LED) was only turned ON to record 1 min during the first 1 min of each NREM and REM post-learning sleep stages. At the end of the session, the mouse was detached from the EEG/EMG recording and moved back to waiting room A where the microscope was removed.

For the optogenetic manipulation experiments (Fig. 6), each mouse in waiting room A was anesthetized immediately after Event 2 using isoflurane for the insertion of two branch-type optical fibers (internal diameter, 0.25 mm) fitted into a housing with a cap, which anchors the inserted optical fiber by screwing around the guide cannula. The tip of the optical fiber protruded 0.2 mm below the guide cannula (DV 1.2 mm from the skull surface). The mouse attached to optic fibers was then moved to its home cage and back to rather a well-lit behavioral room B, placed in a sleep box, and simultaneously connected to the EEG/EMG recording unit and an optical swivel wired to a laser unit (9–12 mW, 589 nm). The delivery of continuous light was manually controlled upon the detection of any sleep stage within 3–4 hours after the CFC and was interrupted whenever the mouse woke up. After the sleep session had ended, the mouse was detached from the EEG/EMG recording and light delivery systems and moved back to waiting room A, where the fiber was removed from the cannula under anesthesia.

Assimilation test (T1): One day after Event 2, one mouse in its home cage was moved from waiting room A into well-lit behavioral room B and then placed in the triangular context (see Extended Data Fig. 1a) for 4 min, except in (Fig. 1a,b) mice were tested in the same pre-exposure context (see Extended Data Fig. 1a,d-f) from Event 1, before being removed, returned to its home cage, and moved back to waiting room A. Contexts were cleaned using water and 70% ethanol after each subject.

Original CFC memory test (T2): One day later, each individual mouse was moved from waiting room A in a blind transport cage to the darkened behavioral room B. The subject was then placed in the conditioned square context (see Extended Data Fig. 1c) with dim lighting and benzaldehyde odor for 4 min, before being removed and moved in the transport cage back to its home cage in waiting room A again. The context was cleaned using water and 70% ethanol after each subject. Circle test (T3): One day later, each mouse in its home cage was moved from waiting room A into well-lit behavioral room B and then placed in the circular context (see Extended Data Fig. 1d) for 4 min before being removed, placed in the home cage, and returned to waiting room A again. The context was cleaned using water and 70% ethanol after each subject.

### Behavioral analysis

All behavioral sessions were conducted using a video tracking system (Muromachi Kikai, Tokyo, Japan) to assess the freezing or motility behavioral patterns of mice. The cumulative duration (s) spent by mice in the complete absence of movement, except for respiration, was considered as the freezing duration. Automated scoring of the freezing response was initiated after 1 s of persistent freezing behavior. Motility assessment was calculated as the cumulative area of movement (pixel size) per 1 min^23^. All sessions were also recorded using Bandicam software (Bandisoft, Seoul, Korea). Freezing behavior of mice with a head-mounted microscope was manually calculated from recorded videos.

### Sleep detection data acquisition and online state detection

All EEG/EMG recordings were made using OpenEx Software Suite (Tucker Davis Technologies, USA), as previously described^41^ but with minor modifications. Briefly, EEG signals were amplified and filtered at 1–40 Hz, while 65–150 Hz was used for EMG; signals were then digitized at a sampling rate 508.6 Hz. Sleep stages were differentiated using an algorithm file that enabled the calculation and analysis of the EMG root mean square value (RMS), EEG delta power (1–4 Hz) RMS, and EEG theta power (6–9 Hz) RMS. The EMG RMS threshold was optimized according to each subject. The mouse was judged to be awake when the RMS exceeded the set threshold value and remained unchanged for three successive 3-s checkpoints. However, when the EMG RMS was lower than the threshold, sleep stage differentiation was concluded based on the delta/theta (d/t) ratio value. Briefly, if the d/t ratio exceeded 1 for the consecutive 9-s checking period, the stage was classified as NREM, and, conversely, it was classified as REM when it was less than 1 for the consecutive 9 s. The state classified by the program was also confirmed by the experimenter through visual inspection of mouse activity and EEG delta-dominant (0.5–4 Hz) or theta-dominant (4–9 Hz) waveforms. EEG/EMG traces recorded during sleep sessions were then extracted using codes written by MATLAB.

### *In vivo* Ca^2+^ imaging data acquisition and analyses

Ca^2+^ signals produced from G-CaMP7 protein^43^ expressed in ACC excitatory neurons were captured at 20 Hz with nVista acquisition software (Inscopix, CA, USA) at the maximum gain and optimal power of LED of nVistaHD. Ca^2+^ imaging movie recordings of all behavioral sessions were then extracted from nVista Data acquisition (DAQ) box (Inscopix, CA, USA). Using Inscopix data processing software (Inscopix, CA, USA), movies were temporally stitched together to create a full movie that contained recordings of all sessions across days, which were spatially downsampled (2x), and then corrected for across-session motion artefacts against a reference frame that was chosen from any session that had a clear landmark “vasculature”. Further motion correction was then applied using Inscopix Mosaic software, as previously described^35, 42^. The full movie was then temporally divided into the individual sessions using Inscopix Mosaic software. Each movie of individual sessions was then low bandpass filtered using Fiji software (NIH) to reduce noise, as described previously^30, 35, 42^. The fluorescence signal intensity change (ΔF/F) for each session was subsequently calculated using Inscopix Mosaic software according to the formula ΔF/F = (F – Fm)/Fm, where F represents each frame’s fluorescence and Fm is the mean fluorescence for the whole session’s movie^30, 35, 42^. Afterwards, movies representing each session were re-concatenated again to generate the full movie for all sessions in the ΔF/F format. Finally, cells were identified using an automatic sorting system, HOTARU^31^, and each cell’s Ca^2+^ signals over time were extracted in a (Ď; time × neuron) matrix format, as previously described^30, 35^. Further processing was performed using codes written in MATLAB to remove the low frequency fluctuation and background noise by subjecting output Ca^2+^ signals to high-pass filtering with a 0.01 Hz threshold, and to then calculate z-scores from the mean of each session, whereby negative values were replaced with zero. Ca^2+^ events were finally extracted after cutting off signals below 3 SD from the local maxima of the ΔF/F signal of each session. Moreover, the mean frequency of each neuron’s Ca^2+^ transients in each session was automatically calculated using another script, also written in MATLAB.

During Event 2 (CFC), cells were classified as Event 2-Shock-responsive (E2-Shock) if they showed 3-fold higher frequencies than the average Ca^2+^ event frequency after the first electric foot shock than that before. Additionally, other cells that were not shock-responsive were classified as either Event 1 (E1)-specific, Event 2 (E2)-Square specific, or non-specific based on their differential Ca^2+^ event frequencies. Briefly, neurons were categorized as Event 1-responsive if they had a 2-fold higher Ca^2+^ spiking frequency in Event 1 relative to the first 2 min of exploration in the square context of Event 2, and vice versa. The remaining neurons were classified as non-specific. The occurrence of Ca^2+^ events of classified neurons that represented each category were automatically enumerated per 500 ms in each session using another MATLAB code. The percent normalized occurrence was then calculated according to the following equation, where (A) is the category of neuron (either E1-specific, E2-Square, or E2-Shock), and *n*(A) is the number of neurons in this category for each mouse:

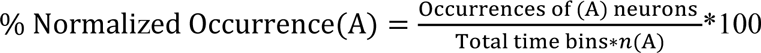

Also, synchronized occurrences – those co-occurring within the same 500-ms time bin – of Ca^2+^ spikes of any of the three classified neuron types were calculated according to the following equation, where (N) represents either or both of the other two neuron categories:

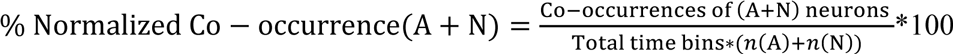

### Histology

At the end of the pharmacological manipulation experiment, mice were deeply anesthetized with 1 ml of combination anesthesia. Using the above-mentioned drug injection system, 0.5 μl of 0.5 μM rhodamine B dye (83689, Sigma-Aldrich) was bilaterally infused into the target brain region immediately before sacrifice and brain extraction for histological analysis. Subjects with misplaced drug injection cannulas were excluded in post-hoc histological examinations. After the optogenetics experiment, mice were deeply anesthetized with 1 ml of combination anesthesia and perfused transcardially with PBS (pH 7.4) followed by 4% paraformaldehyde in PBS (PFA). As previously described^44^, the brains were extracted, then further immersed in PFA for 12–18 h at 4°C. Subsequently, fixed brains were mixed with 25% sucrose in PBS for 36–48 h at 4°C before final storage at –80°C. To obtain coronal sections, brains were sliced into 50 μm sections using a cryostat (Leica CM3050, Leica Biosystems) and then washed in PBS-containing 12-well culture plates (Corning, NY, USA). The sections were further incubated at room temperature for 1 h with a blocking buffer (3% normal donkey serum; S30, Chemicon by EMD Millipore, Billerica, MA, USA) in PBS solution containing 0.2% Triton X-100 and 0.05% Tween 20 (PBST). After the incubation, the buffer was discarded and then rat anti-GFP (04404-84, GF090R, Nacalai Tesque Inc., Japan) primary antibody (1:500) in blocking solution was added for further incubation at 4°C for 24–36 h. At the end of the incubation period, the primary antibody was removed, and sections were washed with 0.2% PBST three times for 10 min each. After washing, sections were treated with a complementary secondary antibody (1:1000), donkey anti-rat IgG Alexa Fluor 488 (A21208, Molecular Probes), in blocking buffer solution at room temperature for 2–3 h. Simultaneously, nuclear staining was performed by adding 1 μg/mL DAPI in the buffer solution (10236276001, Roche Diagnostics). After incubation, treatment was terminated by discarding the solution followed by three 10-min (0.2% PBST) washes before finally mounting the sections on glass slides with ProLong Gold antifade reagent (P36930, Invitrogen, USA).

### Confocal microscopy

Images were acquired using a confocal microscope (Zeiss LSM 780, Carl Zeiss, Jena, Germany) with a 0.8 numerical aperture, 20× Plan-apochromat objective lens. All parameters were standardized within each magnification and each experimental condition.

### Statistics

Statistical analyses were performed using Prism 6.01 (GraphPad Software, San Diego, CA, USA). Unless otherwise specified, comparisons between multiple groups were made using an ANOVA with post-hoc tests, as shown in the corresponding figure legends.

## Data and code availability

The data and codes that supported the findings of this study are available from the corresponding author upon reasonable request.

## Acknowledgments

We are grateful to K. Deisseroth (Stanford University) for providing eArch3.0-eYFP cDNA; H. Bito (University of Tokyo) for the E-SARE cDNA; N. Ohkawa for hKikGR cDNA; A. Konno and H. Hirai for disclosing the AAV_9_ virus production protocol prior to publishing; and S. Okami for maintaining the mice. We thank K. Choko, M.E. Wally, A. Suzuki, K. Ghandour, E. Murayama, S. Nakayama, N. Oishi (University of Toyama), Y. Saitoh (Dokkyo Medical University), M.J. Alam (University of Michigan), and J. Yokose (University of Texas Southwestern Medical Center) for technical teaching and support. All members of the Inokuchi laboratory supported and discussed this study. This work was supported by JSPS KAKENHI (grant number JP18H05213), the Core Research for Evolutional Science and Technology (CREST) program (JPMJCR13W1) of the Japan Science and Technology Agency (JST), a Grant-in-Aid for Scientific Research on Innovative Areas “Memory dynamism” (JP25115002) from MEXT, and the Takeda Science Foundation (to K.I.) and by the Uehara Memorial Foundation (to M.A.). Additional support was kindly provided by a Grant-in-Aid for young scientists from JSPS KAKENHI (JP19K16892) (to K.A.), JSPS KAKENHI (JP20H03554 and JP17K19445), THE HOKURIKU BANK Grant-in-Aid for Young Scientists, the FIRSTBANK OF TOYAMA SCHOLARSHIP FOUNDATION RESEARCH GRANT, and the Takeda Science Foundation support (to M.N.).

## Author contributions

M.A., K.A., and K.I. designed the experiments and wrote the manuscript. M.A., K.A., and R.O-S. performed the experiments. M.A. and K.A. analyzed the data. M.N. generated MATLAB codes. K.I. supervised the entire project.

## Competing interests

The authors declare no competing interests.

**Correspondence and requests for materials** should be addressed to K.I.

---

**Extended Data Fig. 1.**
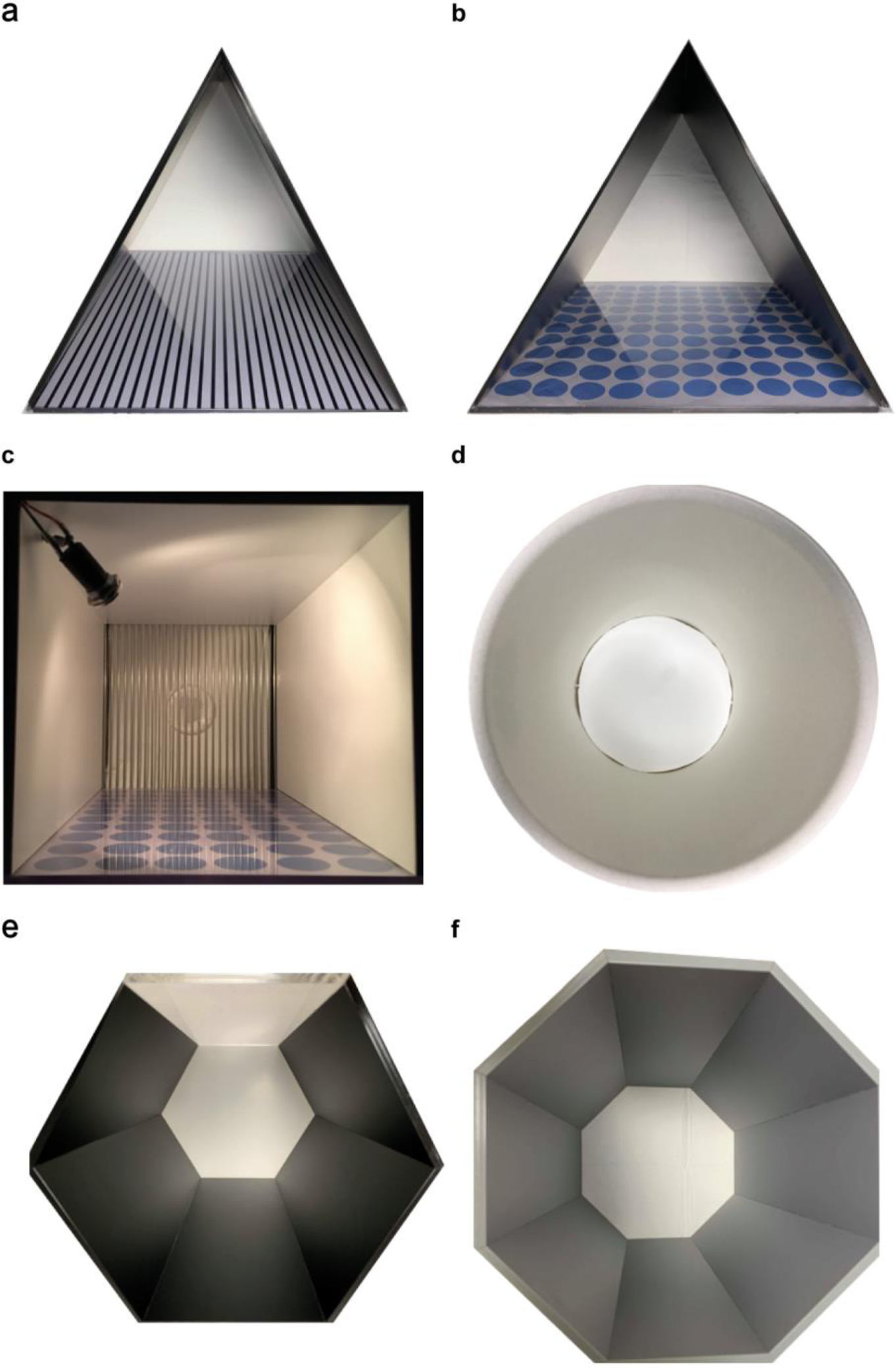
Top view of the contexts used in the study. **a,b,** Triangle-shaped prism context with two black walls, a plexiglass front covered from the outside with a white piece of paper showing either a black striped **(a)** or a blue circled **(b)** pattern, and a transparent acrylic floor placed on a white paper towel (Kim Towel, Kimberly-Clark, USA) (one side × height: 290 × 320 mm). **c,** Square-shaped cubic context (width × depth × height: 175 × 165 × 300 mm) with three off-white sides, a plexiglass front covered from outside with a white piece of paper showing a blue circled pattern, and a flooring of 26 stainless steel rods with a diameter of 2 mm placed 5 mm apart. **d,** Circle-shaped cylindrical context (diameter × height: 205 × 320 mm) with off-white walls and a white acrylic floor. **e,** Hexagon-shaped context (one side × height: 100 × 300 mm) with five black walls, a plexiglass front covered from the outside with a white piece of paper, and a transparent acrylic floor placed on a white paper towel (Kim Towel, Kimberly-Clark, USA). **f,** Octagon-shaped context with eight gray walls (one side × height: 70 × 300 mm) and a white acrylic floor. All contexts were placed in exactly the same location during behavioral experiments.

**Extended Data Fig. 2.**
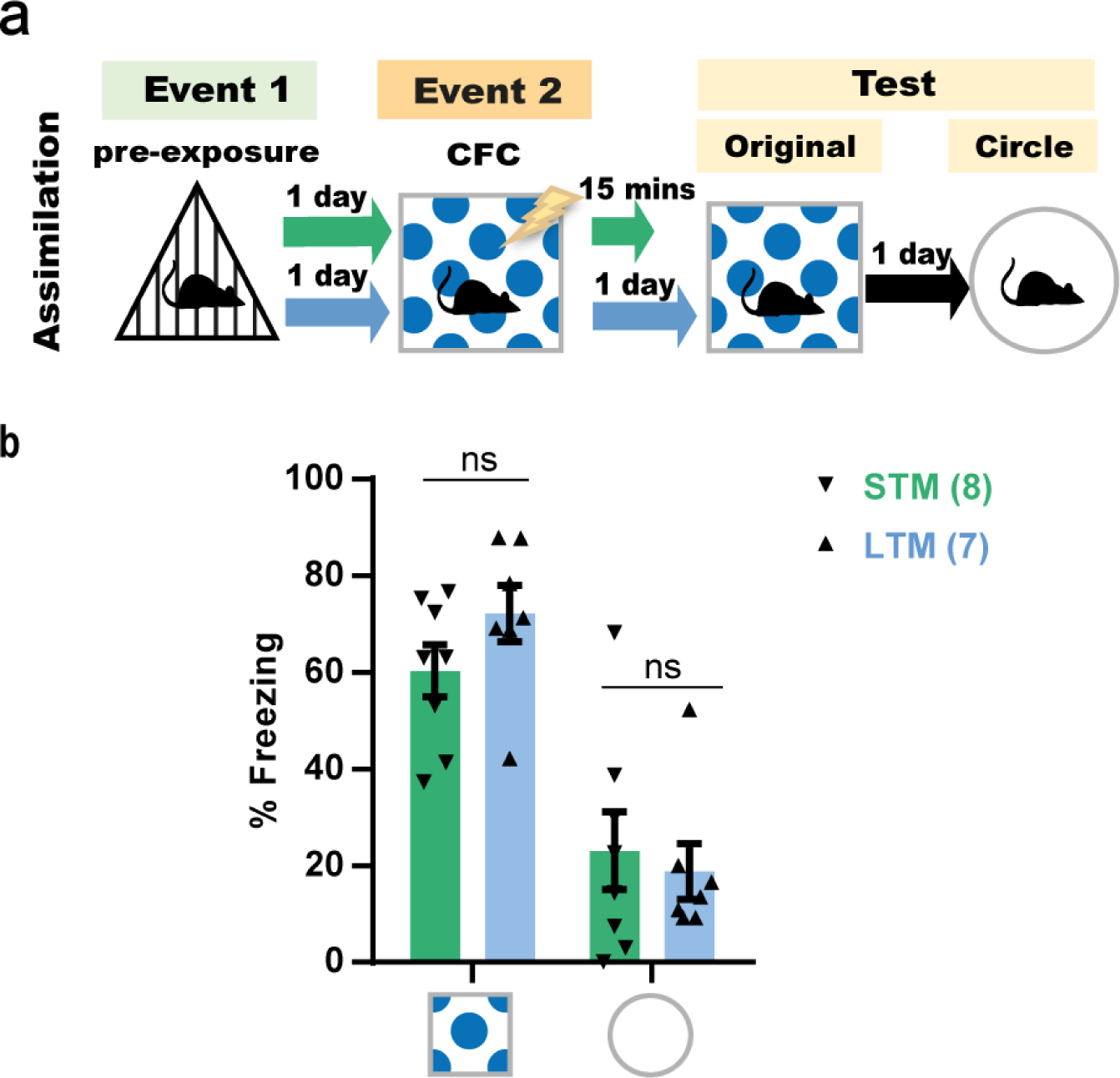
The original CFC memory shows no significant incubation-dependent difference in freezing between short-term and long-term testing intervals. **a,** Experimental design. **b,** Freezing levels during test (4 min). *P*-values were determined using a two-way RM ANOVA followed by Sidak’s multiple comparisons. Data are presented as mean ± SEM. ns, not significant (*P* > 0.05).

**Extended Data Fig. 3.**
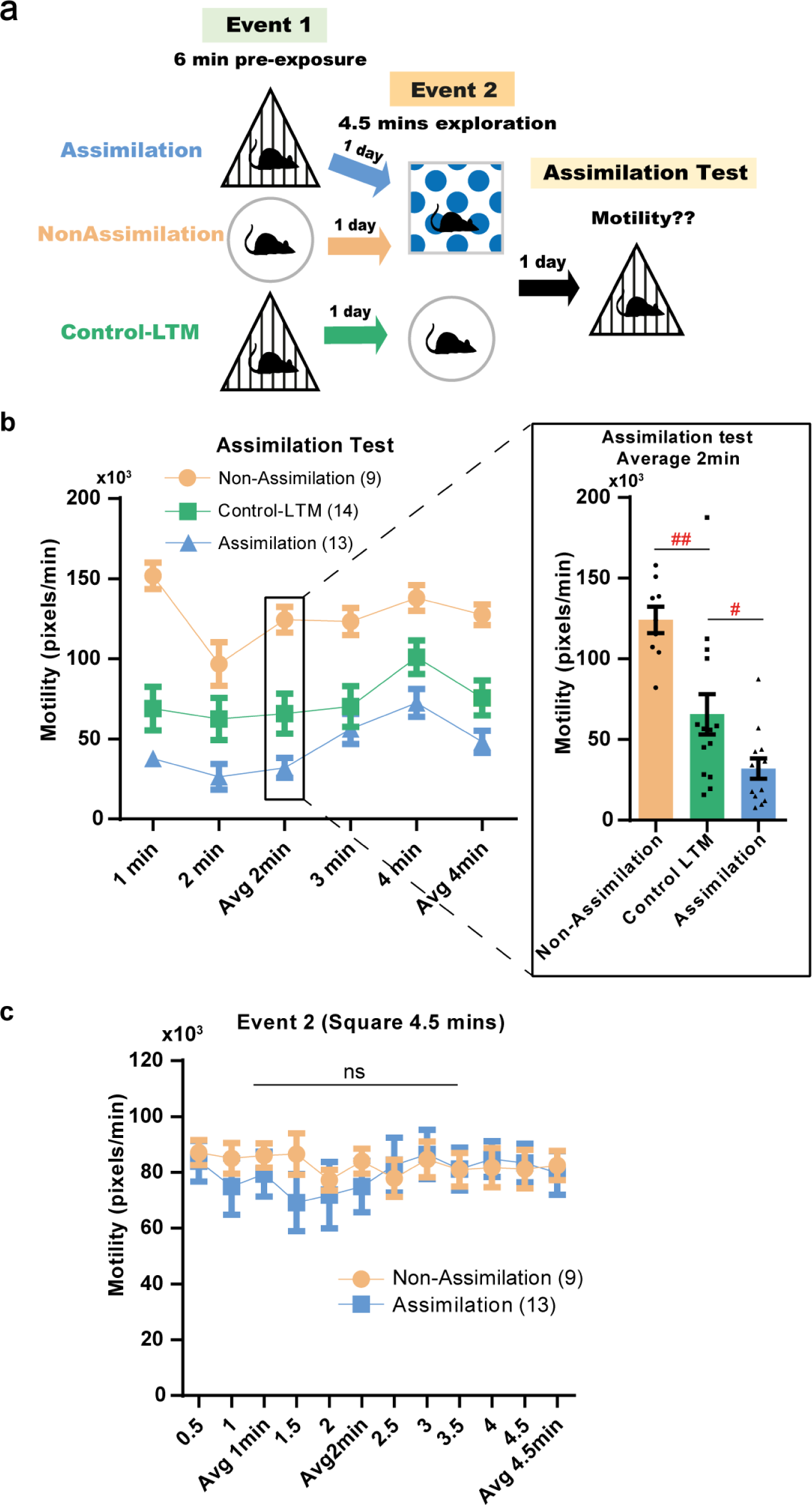
Pre-exposure to spatial commonalities induces offline assimilation of non-emotional memories. **a,** Experimental design. **b,** Motility examined in 1-min intervals during the test (4 min) in the triangle context. Inset, average motility calculated for the first 2 min of the test. *P*-values were determined using a one-way ANOVA followed by Tukey’s multiple comparisons, ^#^*P* < 0.05, ^##^*P* < 0.01. Data are presented as mean ± SEM. **c,** Motility was measured in 0.5-min intervals during Event 2 (4.5 min) in the square context for the assimilation and non-assimilation groups. *P*-values were determined using a two-way RM ANOVA followed by Sidak’s multiple comparisons. ns, not significant (*P* > 0.05). Data are presented as mean ± SEM.

**Extended Data Fig. 4.**
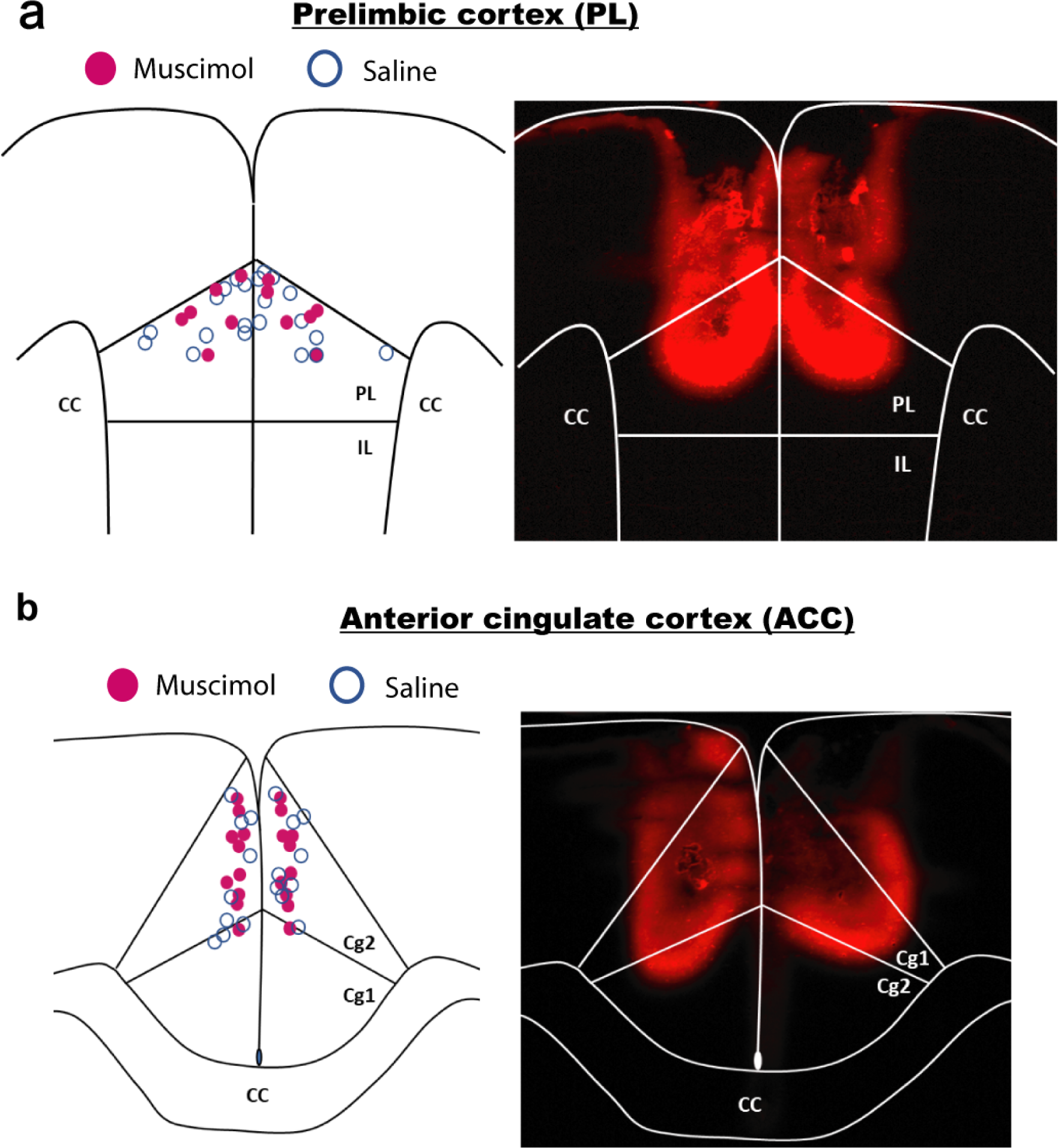
Drug injections into the ACC or PL. **a,** Left: Schematic diagram showing the center of each injection in the PL. Right: Schematic diagram showing representative rhodamine B staining in the PL. **b,** Left: Schematic diagram showing the center of each injection in the ACC. Right: Schematic diagram showing representative rhodamine B staining in the ACC. IL, infralimbic cortex; CC, corpus callosum.

**Extended Data Fig. 5.**
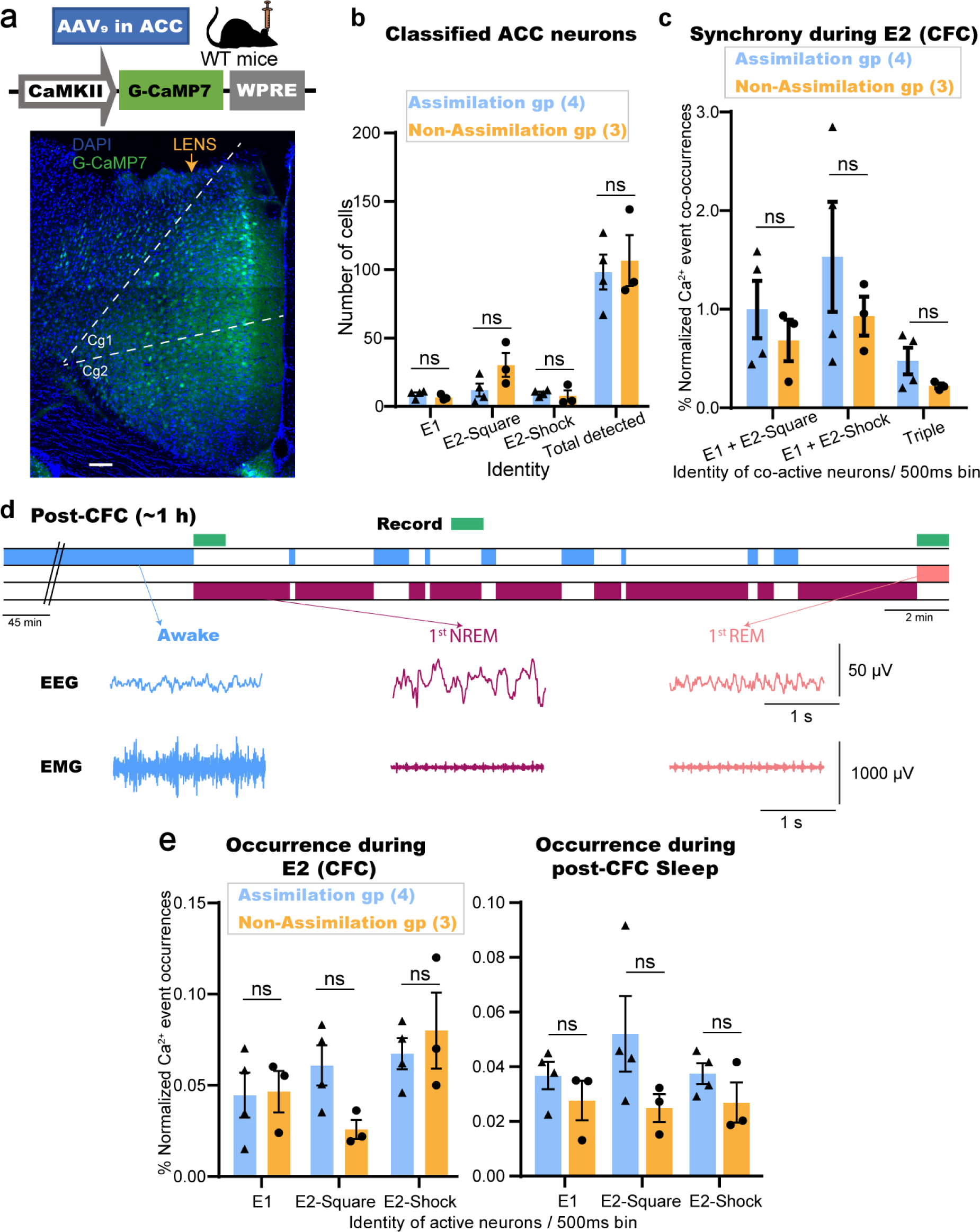
*In vivo* calcium imaging during offline memory assimilation task as well as sleep detection during post-CFC offline session. **a,** Top: Design of AAV injected into the ACC. Bottom: Coronal section of the ACC showing G-CaMP7 protein expression and GRIN lens trace. Scale bar represents 100 μm. **b,** Averaged number of ACC cells classified as E1, E2-Square, or E2-Shock-responsive as well as mean total detected in each group. **c,** % Normalized Ca^2+^ event co-occurrences of E1-cells and E2-cells throughout the 2.5 min post-shock period in CFC training. **d,** Top: Diagram for post-CFC stage-specific recording. Bottom: Example EEG and EMG recordings for each detected stage. **e,** Single Ca^2+^ event occurrences of E1- or E2-responsive cells throughout the 2.5 min post-shock period in CFC training (**e**, left), or within 2-min post-CFC sleep (**e**, right). *P*-values were determined using unpaired *t* test (**b and e**) and Wilcoxon rank-sum test **(c)**. ns, not significant (*P* > 0.05). Data are presented as Mean ± SEM in (**b, c, and e**).

